# Parasitic Infection by *Pseudocapillaria tomentosa* is Associated with a Longitudinal Restructuring of the Zebrafish Gut Microbiome

**DOI:** 10.1101/076596

**Authors:** Christopher A. Gaulke, Mauricio L Martins, Virginia Watral, Michael L. Kent, Thomas J. Sharpton

**Author notes:** Corresponding Author: Thomas J Sharpton, Department of Microbiology and Department of Statistics Oregon State University, 97330.

## Abstract

Helminth parasites represent a significant threat to wild, domesticated, and research animal health. *Pseudocapillaria tomentosa* is a common intestinal nematode parasite and an important source of infection in zebrafish. Symptoms of the infection vary widely from no clinical signs to sever emaciation and mortality, however, the reasons underpinning these disparate outcomes are unclear. Components of the microbiome may interact with parasites to influence their success in the gut while parasite infections are also known to influence the composition of the gut microbiome. In this study we evaluated the longitudinal changes in the gut microbiome structure and gut physiology during experimental *P. tomentosa* infection in adult 5D line zebrafish. We observed less severe signs of infection and less mortality in these fish than previously described in AB line fish. However, inflammation and epithelial hyperplasia in the intestine was still observed in infected 5D line fish. The composition of the microbiome changed rapidly during the infection and these changes were associated with parasite stage of development and burden. Individual taxa covaried with parasite abundance in the intestine intimating the gut microbiome may influence parasite burden. Associations between taxa and parasite abundance in some cases appeared to be phylogenetically patterned. Strong positive associations were observed between OTUs phylotyped to Proteobacteria and abundance of adult parasites and parasite eggs. Together these experiments demonstrate that *P. tomentosa* infection results in a rapid and temporally dynamic disruption of the zebrafish gut microbiome and clarify how interactions between the gut microbiome and intestinal parasites may impact fish populations.

## Introduction

Wild animals are frequently exposed to and infected by intestinal parasites [1,2]. While a relatively small number of individuals in a population are infected at levels that result in mortality, many will carry parasitic loads that influence host growth [3], behavior [4], or reproductive fitness [5]. As a result, parasitic infections can act as a significant selective force on a population [6]. Often, the specific factors that determine a parasite's success in the gut [7] and the mechanisms through which infection impacts host physiology are not well described[8]. Efforts to determine how animals change over the course of parasitic infection are useful for discerning these properties[9], which ultimately require elucidation to ensure effective management and preservation of animal populations and understand their evolutionary dynamics.

A growing body of evidence indicates that the gut microbiome may interact with intestinal parasites to influence their growth or mediate their physiological effect on the host. For example, helminth-infected humans [10,11] and mice [12] harbor gut microbiomes with significantly different structures and diversity than uninfected controls. Additionally, studies in mice have found that specific gut bacteria are perturbed upon helminth infection to yield alterations in host immune status [13]. Other work has shown that the gut microbiome acts as an innate immune barrier to intestinal infection [14], intimating that specific bacteria may attenuate parasitic infection. However, there is limited insight into how the developmental variation of parasitic populations within the gut, which may include multiple life history stages including maturation and reproduction, associates with the structure and diversity of the gut microbiome. Monitoring the variation of the gut microbiome over the course of infection may clarify which microbiota influence or are impacted by the development of intestinal parasitic populations.

Here, we use a zebrafish (*Danio rerio*) model to clarify the longitudinal co-variation between intestinal parasitic infection and the gut microbiome. *Pseudocapillaria tomentosa*, a capillarid nematode that preferentially infects the guts of fish, is an important cause of disease in zebrafish facilities [15]. The lifecycle of *P. tomentosa* can be direct or indirect utilizing oligochaetes as a paratenic host [16]. In the gut, *P. tomentosa* causes intestinal inflammation, tissue damage and epithelial hyperplasia [17] and fish infected with *P. tomentosa* often appear emaciated and lethargic, though cryptic subclinical infections have also been reported [15]. We monitored how *P. tomentosa* infection associates with the zebrafish gut microbiome over the course of infection. We find that early time points of infection were associated with mild to moderate inflammation that increased over time, consistent with prior studies [17]. *P. tomentosa* infection also associates with an altered microbial community composition in a parasite life-history stage dependent fashion. Additionally, parasitic burden at various life-history stages is correlated with the abundance of specific microbiota, intimating their interaction. Our study clarifies how fish, their gut microbiota, and intestinal parasites interact and intimates that the gut microbiome may be an important factor in the population-level dynamics of parasitic infection.

## Methods

### Parasite infection and burden quantification

The use of zebrafish in this study was approved by the Institutional Animal Care and Use Committee at Oregon State University (permit number: 4800). To create an infectious environment 30 *P. tomentosa* infected (donor) zebrafish were placed in an 80 L static flow tank for several weeks. Donor fish infection was confirmed by examining the feces for the presence of *P. tomentosa* eggs using light microscopy. Prior to the initiation of the experiment these fish sequestered in a net cage that was perforated such that feces from infected fish would pass into the tank below, maintaining an infectious environment, while physically isolating these fish from the bottom portion of the tank. To infect fish, 65 *P. tomentosa* naïve adult 5D line zebrafish (recipient fish) were placed in the exposure tank for three days (**Figure 1**). After exposure the recipient fish were removed and randomly separated into six 3 L tanks (n=10) and one additional 3 L tank (n=5). Fish from these tanks were progressively evaluated at 6, 11, 18, 25, 32, 39 and 46 days post-initial exposure (p.e.). Seventy-two hours before necropsy fish were isolated and individually housed in 1.5L tanks for fecal collection. All feces present was collected from each tank every 24hrs during the 72hr period and the last sample (72hrs post isolation) was stored at - 20°C until processing. During the period of feces collection the water quality were kept at: temperature 27.60±0.80°C, pH 7.50±0.20, total ammonia 0.19±0.05 mg **·** L^-1^ measured with a colorimetric kit (Aquarium Pharmaceuticals, Inc.) and dissolved oxygen 6.59±0.29 mg**·**L^-1^ measured with an oximeter (Fisher Scientific, Texas). After the fecal collection the fish were euthanized by immersion in cold water, and the intestines were removed for parasitological analysis of the intestines. Wet mounts were prepared from each intestine and examined with light microscopy to quantify the number of eggs, larvae, and adult worms present.

**Figure 1.**
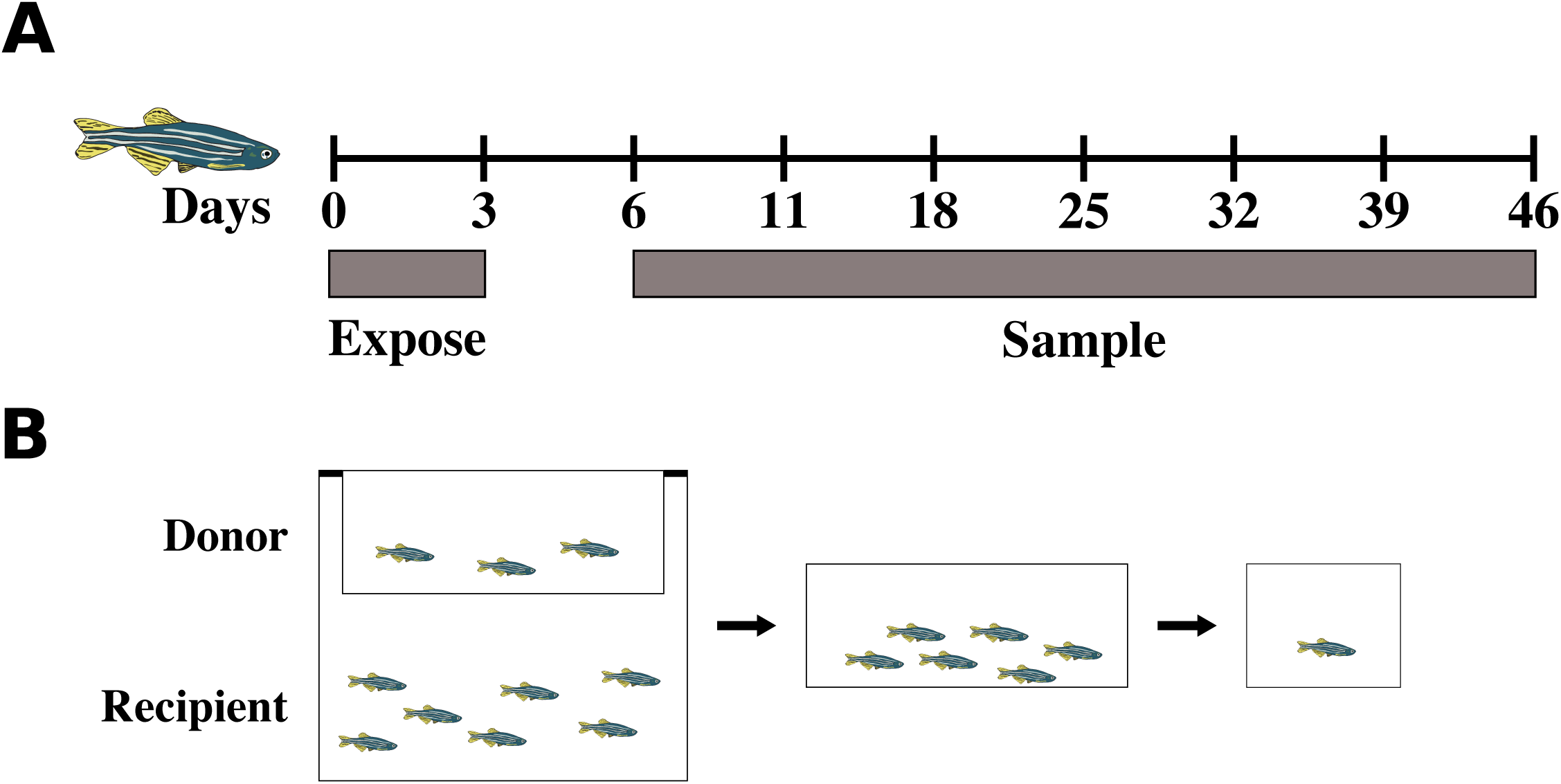
Experimental timeline. Sixty-five 5D line zebrafish were exposed to *Pseudocapillaria tomentosa* infected zebrafish. After exposure fish were transferred to 3 L tanks and serially evaluated at seven time points. **A)** An experimental timeline of exposure and sample collection. **B)** Experimental set up of exposure and sample collection.

### Intestinal Histology

We exposed an additional, parallel group of fish to the parasite to elucidate the pathological changes associated with the development and microbiome changes. Here 30 fish were exposed as above, and sampled at 8, 15, 28, and 42 d post initial exposure. Fish were euthanized, preserved in Dietrich’s fixative, processed for histology, and stained with hematoxylin and eosin using our standard protocol [18].

### 16S amplicon library preparation and sequencing

Isolation of microbial DNA from fecal samples was performed using the MoBio PowerSoil^®^ DNA isolation kit (MOBIO, Carlsbad, CA USA) following the manufacturer’s protocol. An additional 10-minute incubation at 65˚C before bead beating was added to facilitate cellular lysis. Immediately following this incubation the samples underwent bead beating on the highest setting for 20-minute using Vortex Genie 2 (Fisher, Hampton, NH USA) and a 24-sample vortex adaptor (MOBIO). One microliter of DNA was then used as input into triplicate PCR reactions and the remaining DNA stored at −20˚C. Amplification of the V4 region of the 16S rRNA was performed as previously described [19,20]. To ensure proper amplification, amplicons were visualized using gel electrophoresis and quantified using the Qubit^®^ HS kit (Life Technologies, Carlsbad, CA USA) according to the manufacturer’s instructions. A total of 200ng of amplicon library was pooled and the pooled library was then cleaned using the UltraClean^®^ PCR clean-up kit (MOBIO) and diluted to a concentration of 10nM. The final pooled and cleaned product was submitted to the Oregon State University Center for Genome Research and Biocomputing (CGRB) for cluster generation and 250bp paired end sequencing on an Illumina MiSeq instrument. This generated ~1.3 million 250bp paired end which were input into QIIME [21] for open reference OTU picking and taxonomic assignment using the UCLUST [22] algorithm and a 97% identity threshold against the Greengenes (version 13_8) reference [23].

### Statistical analysis

Statistical analysis was conducted on a QIIME generated rarefied BIOM table (sampling depth 8,000 counts) in R. The dataset was first filtered to remove OTUs present in fewer than ~10% of the samples. The resulting filtered dataset, which consisted of 785 OTUs, was used for downstream analysis. Kruskal-Wallis tests with a pairwise Mann-Whitney U post-hoc tests (false discovery rate p-value correction) were used to determine phylotypes that significantly differed across time points.

Beta-diversity was measured using Bray-Curtis distance, and non-metric multidimensional scaling (NMDS) was used to quantify and visualize compositional similarity of communities. Significant differences in beta-diversity associated with parasite burden were calculated using Permutational Multivariate Analysis of Variance (PERMANOVA, vegan::adonis) with 5,000 permutations.

Spearman’s rank correlation coefficients were calculated for OTU abundance and parasite burden parameters including number of eggs, adults and larvae present in the intestine. Significant and moderate to strong correlations (|rho| ≥ 0.4, fdr < 0.05) were then subjected to linear modeling of OTU abundance by modeling abundance vs. time and abundance vs. time plus burden parameters. OTUs for which the inclusion of burden significantly increased fit (analysis of variance; p <0.05, fdr < 0.05) were retained. A heatmap was generated using the OTUs that passed filtering in R (gplots::heatmap.2) using unsupervised hierarchical clustering with default clustering parameters. The spearman’s correlation coefficients were used in this visualization.

Permutation tests (100 permutations) were used to determine if clustering of OTUs associated with specific classes of bacteria was random across the major bifurcations of the heatmap dendrogram. We restricted this analysis to the first two bifurcations of the dendrogram such that six subdendrograms that represented unique association patterns with parasite burden parameters were produced. Only the three most abundantly represented classes (Gammaproteobacteria, Betaproteobacteria, and Fusobacteria) were considered in this analysis. The number of OTUs associated with each class was tabulated for each subdendrogram. For each permutation the class label was randomly assigned to a tip on the dendrogram and the number of classes corresponding the subdendrogram was tabulated. A one-tailed z-test was then used to determine if a subtree, or branch of the dendrogram, contained more members of a certain class than expected if the classes were distributed randomly. False discovery rate was controlled at fdr < 0.05 (stats::p.adjust).

## Results

### *Parasite infection in* Danio rerio

To determine the impact of *P. tomentosa* infection on the gut microbiome of zebrafish, we followed the progressive impacts of the infection on the microbiome in 65 adult zebrafish. Briefly, adult zebrafish were infected with *P. tomentosa* by exposing them to the feces of actively infected zebrafish for 3 days (**Figure 1**). The earliest time point examined (6d p.e.) after exposure to the parasite showed no evidence of parasite burden (**Figure 2; Sup Figure 1**). Larvae were first observed 11d p.e., which coincided with peak larval parasite burden in these animals. Larval burden decreased rapidly after 11d p.e. and larvae were absent completely from all fish by 46d post exposure. Shortly after the presence of larvae was detected, adult worms were first observed (18d p.e.). Adult worm burden peaked at 18d p.e. and then declined until the final time point (**Figure 2; Sup Figure 1**). Following the appearance of adults, eggs were observed beginning at 25d post exposure. Presence of eggs indicated presence of sexually mature worms, and eggs counts represent both eggs free in the lumen and within female worms. Mean intestinal egg abundance peaked at 39d p.e. and then declined at day 46 (**Figure 2; Sup Figure 1**). The prevalence of *P. tomentosa* infection in fish after 6d p.e. was 100% at all time points with the exception of 39d p.e., where we observed parasite burden in 9 of the 10 fish.

**Figure 2.**
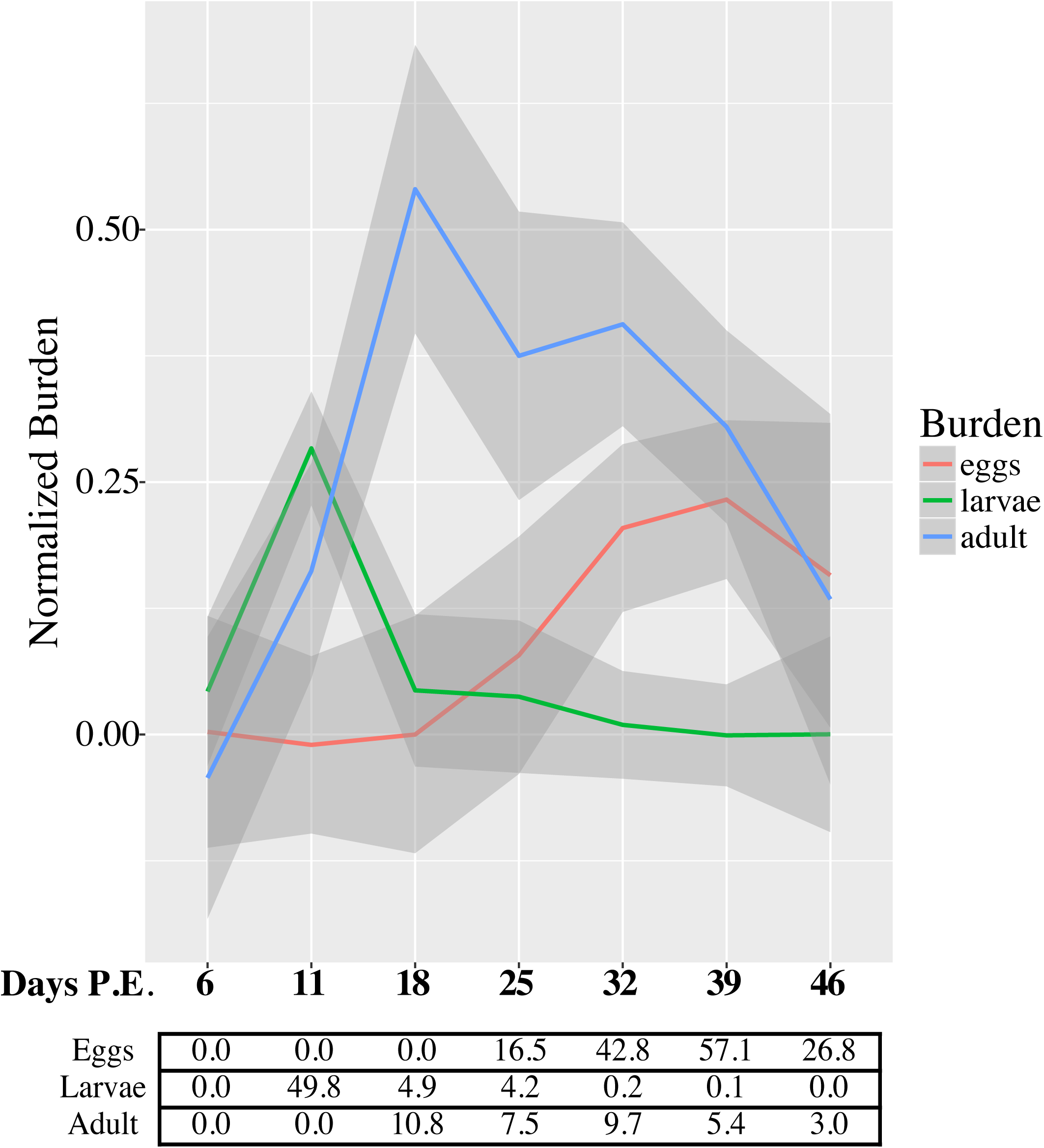
Intestinal parasite burden across the length of infection. The colored lines indicate normalized abundance (mean abundance/ max abundance) of egg, larval, and adult parasite burden across the length of infection. Ribbons surrounding lines indicate the standard error of the mean at each time point. The table of values below the graph contains the mean abundance of each parameter at the indicated time point.

Interestingly, there was very low mortality during the experiment (1/65) suggesting that 5D line zebrafish might be more robust in the face of *P. tomentosa* infection than other, more susceptible lines that have high mortality rates upon infection [24].

During quantification of parasite burden it was necessary to crush the intestine to rapidly making histological investigations of the impacts of parasite burden difficult. Therefore, a separate cohort of fish exposed to the *P. tomentosa* for pathology analysis (**Figure 3**). Fish were infected as above and followed for 42 days after infection. As with the fish used for microbiome investigation the fish in this cohort appeared clinically normal. Worms were only detected in the epithelium and lumen and pathological changes were confined to the lamina propria and epithelium. Early in the infection (e.g. 8d p.e.), structures consistent with necrotic or apoptotic cells were frequently observed throughout the epithelium (**Figure 3A, B**). At 15 d p.e. worms were larger, but no eggs were observed within worms (**Figure 3C**). Whereas confined to the epithelium, the underlying lamina propria exhibited mild to moderate chronic inflammation. The extent of the inflammatory response increased through the remainder of the experiment (**Figure 3D-F**), and fish from the last sample (42 d p.e.) also exhibited hyperplasia of the epithelium.

**Figure 3.**
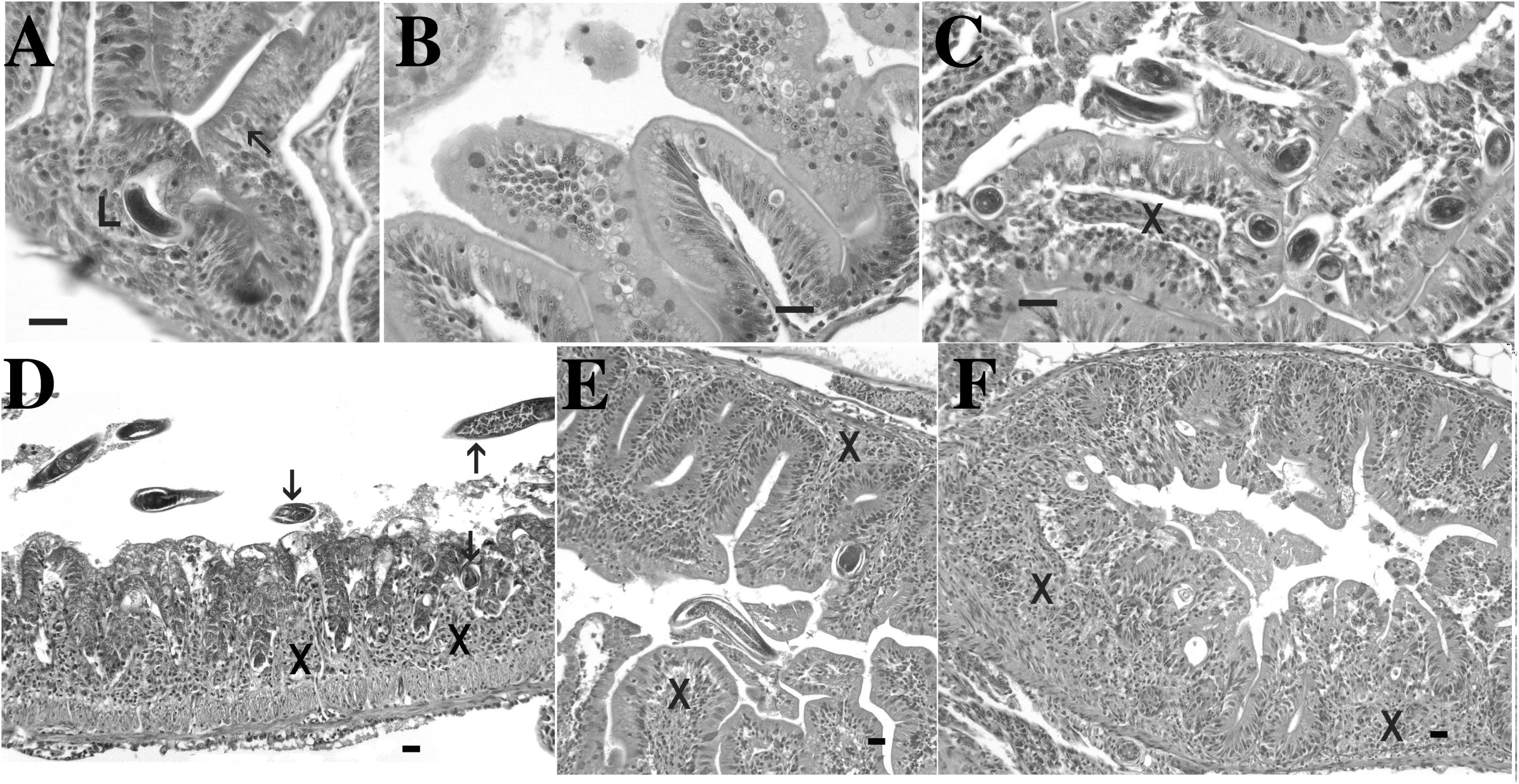
Progression of infection and associated lesions in the intestines of zebrafish experimentally infected with *Pseudocapillaria tomentosa*. Hematoxylin and eosin stained sections of zebrafish intestine. Bar = 20 µm. **A, B)** 8 d post-exposure. L = larval worm in epithelium. Arrows = necrotic/apoptotic bodies in epithelium. **C)** 15 d p.e. nematodes in epthelium with mild, chronic inflammation (arrows) in lamina propria. **D)** 28 d p.e. prominent, diffuse, chronic inflammation in lamina propria (X) with nematodes (arrows) free in the lumen and in epithelium. **E, F)** 42 d p.e. prominent, diffuse, chronic inflammation in lamina propria (X) with nematodes (arrows) free in the lumen and in epithelium. Infection is also associated with diffuse epithelial hyperplasia.

### Pseudocapillaria tomentosa infection is associated with rapid restructuring of the microbiome

To determine if *P. tomentosa* infection resulted in changes in microbiome structure, we examined the taxonomic composition of the microbiome using 16S amplicon sequencing across the length of the experiment. Regardless of length of time post exposure the zebrafish microbiome was dominated by two phyla, Proteobacteria and Fusobacteria consistent with previous studies in fish [25,26]. The phyla Bacteroidetes and Proteobacteria were core across all fish (i.e., present in 100% of samples). Other phyla including Fusobacteria, Tenericutes, Firmicutes, and Cyanobacteria were also highly prevalent in these fish (present in > 50% of samples). The abundance of the phylum Bacteroidetes increased in *P. tomentosa* infected fish from 11d p.e. (p < 0.05) when compared to fish that did not have parasite burden (i.e., 6d p.e.). Fusobacteria abundance significantly increased at 11d p.e. (p < 0.05), while Proteobacteria abundance was significantly decreased (**Figure 4A**; p < 0.05). These changes correspond to the first signs of parasite burden in this cohort (i.e., first observation of larvae). The abundance of Tenericutes was also in fish with parasite burden when compared to fish from the 6d p.e. group. Infection with *P. tomentosa* also associated with altered community structure of the zebrafish gut microbiome (**Figure 4B**). Permutational Multivariate Analysis of Variance (PERMANOVA) indicated that microbial communities were significantly stratified by time post exposure as well as intestinal eggs (p < 0.05), larvae (p < 0.0005), and adult worms (p < 0.005) abundance (**Figure 4B-E)**. Intragroup beta-diversity (Bray-Curtis) was decreased in all groups with parasite burden except the 46d p.e. group, which also had lowest parasite burden of any group after 6d p.e., when compared to 6d p.e. fish indicating that the fish from these groups are more homogenous in their microbiome composition (**Sup figure 2**). Interestingly, intergroup variability between 6d p.e. and 46d p.e. was significantly elevated when compared to those between 6d p.e. and the other time points (**Sup figure 2**). This indicates that while individuals within these groups may be more diverse when compared to other individuals in the same group, individuals in these two groups are more dissimilar in terms of composition than 6d p.e. group fish are to any other group of fish (**Sup figure 2**). This did not appear to be due to changes in alpha diversity as no significant differences in alpha diversity were observed across the groups. Together these data indicate that infection with *P. tomentosa* is associated with altered gut microbial community structure.

**Figure 4.**
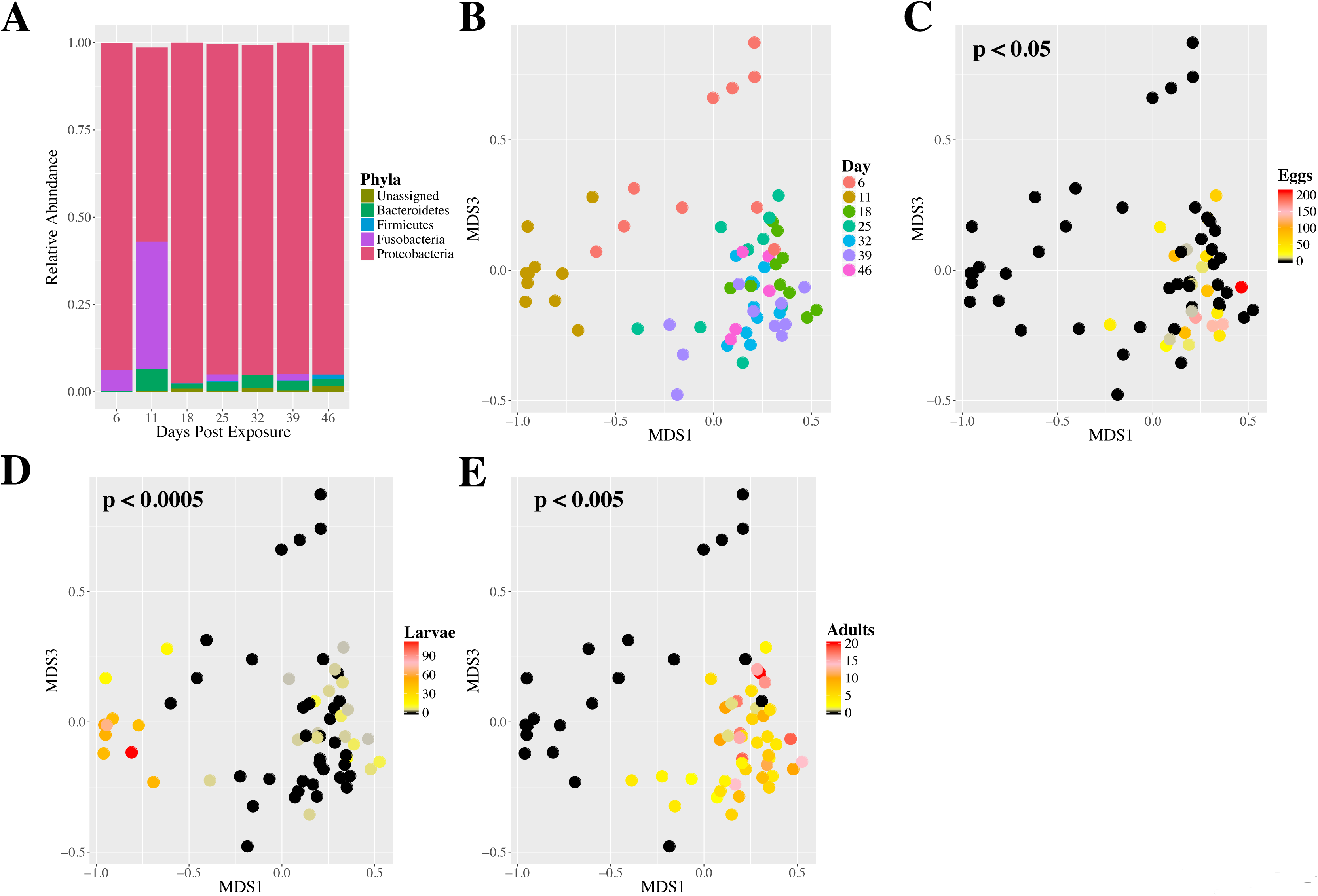
*Pseudocapillaria tomentosa* infection rapidly alters the microbiome of zebrafish. **A)**Abundance of the five most highly abundant phyla across all fish at each time point. **B-E)**
Non-metric multidimensional scaling plot of fish exposed to *P. tomentosa*. Dots are colored by **B)** the days post exposure, **C)** abundance of eggs, **D)** abundance of larvae, **E)** abundance of adults. PERMANOVA test were conducted to determine if variance across different parameters was significant and the p value is indicated for each graph.

**Figure 5.**
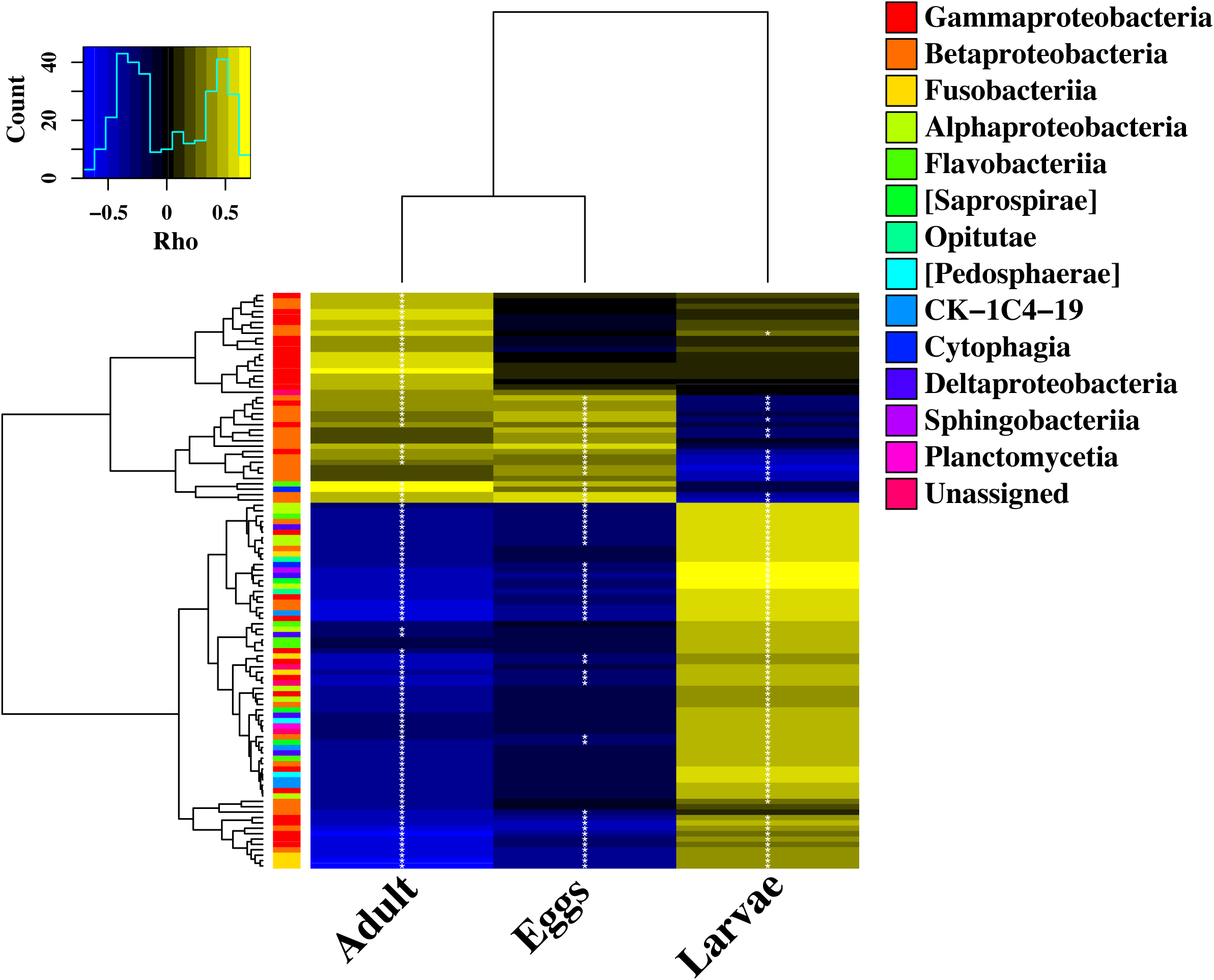
Gut microbes are associated with *P. tomentosa* burden. A heat map of correlations between OTU abundances and parasite burden parameters. Yellow color indicates positive correlations, blue color indicates negative correlations. Asterisks indicate significant correlations (fdr < 0.05). Color bar on the left side of graph indicates the class of each OTU. Dendrograms represent result of unsupervised hierarchical clustering.

### Changes in microbiome structure are correlated with Pseudocapillaria burden

Interactions between helminths and the microbiome can facilitate, or disrupt the ability of a parasite to colonize the host [14]. However, little is known about the potential interactions between *P. tomentosa* and microbiome composition in fish. To identify potential microbe-parasite interactions we calculated Spearman’s correlation coefficients between OTU abundance and abundance of eggs, adults, and larvae in the intestine. Moderate-to-strong statistically significant correlations (|rho| > 0.4; fdr < 0.05) were selected for downstream analysis. As the parasite burden may also be correlated with time, each of OTU-burden pairs were then subjected to two regression analyses. First OTU abundance was regressed against the time, then OTU abundance was regressed against time plus burden. Analysis of variance (ANOVA) was then used to compare these two models and the only models that incorporated burden, improved the model fit (R^2^), and reached the significance threshold (p < 0.05) were retained. These pairs were used to build a heat map of spearman correlations across all burden parameters (**Figure 4**). A cluster of 36 OTUs were positively correlated with the number of adult worms present per fish. The OTUs in this cluster were largely associated with the phyla Proteobacteria (33 of 36), and included OTUs associated with the genera *Acinetobacter*, *Vogesella*, and the families Aeromonadaceae, Oxalobacteraceae, Comamonadaceae, and Neisseriaceae. Interestingly, members of the family *Acinetobacter* are known to be opportunistic pathogens of fish [27]. Another small cluster of 13 OTUs were negatively correlated with adults and eggs while being positively associated with larvae. These OTUs were associated with the phyla Proteobacteria (10 of 13) and Fusobacteria (3 of 13). All of the OTUs associated with the phylum Fusobacteria were associated with the genus *Cetobacterium*, a genus present in the gastrointestinal tracts of many warm water fishes [28]. Decreased *Cetobacterium* has been observed under starvation in zebrafish [29] and might reflect nutrient limited conditions in the gut that are common with intestinal parasites [3]. Together, these data indicate that infection with *P. tomentosa* is associated with shifts in the zebrafish gut microbiome and that some of these changes are correlated with parasite burden parameters.

Given the clustering patterns observed across correlation coefficients we next asked if these clusters were statistically enriched for specific classes of microbes. We restricted this analysis to the three most abundant classes (Betaproteobacteria, Gammaproteobacteria, and Fusobacteriia) in the OTU heat map and the first two bifurcations of dendrogram, which included four distinct and common patterns of association with different *P. tomentosa* burden (Figure 4). In total this created six clades with unique patterns of association with *P. tomentosa*: 1) clade 1 harbored taxa that were largely positively correlated with adults worms, and had weak or no association with eggs and larvae, 2) clade 2 taxa were positive correlated with adults and eggs and negatively with larvae, 3) clade 3 contained clades one and two, 4) clade 4 taxa were generally exhibited strong positive associations with larvae and negative associations with adults and may or may not be negatively associated eggs, 5) taxa in clade 5 taxa had strong negative assocaitions with adults and eggs and moderate positive associations with larvae, 6) clade 6 contained clades 4 and 5. We used permutation tests to determine if any of these clades were significantly enriched for a specific class or classes. Gammaproteobacteria was enriched in clade 1 and 3, Betaproteobacteria was enriched in clade 2 and 3, and Fusobacteriia was significantly enriched in clade 5. Parasitic helminth infection is known to increase the ability of some Proteobacteria to colonize the gut and cause disease [30]. It is also possible that the bacteria themselves promote parasite burden. The data suggest phylogenetic patterns might exist in how microbiomes change in response to infection with *P. tomentosa*. Further, these data are also consistent with the hypothesis that specific classes of microbes might influence the infectivity or fecundity of *P. tomentosa*.

## Discussion

A growing body of evidence suggests that the gastrointestinal microbiome performs vital roles in maintenance of host health and homeostasis [31–33]. For example, the microbiome contributes to digestion, growth, and immune function in fish [34–36] the latter of which includes the fish microbiome's action as an innate immune barrier. The fish microbiome is shaped by several factors including developmental stage, [37,38], chemical exposures [25], and diet [36]. To date, it is unclear how infection by intestinal parasites impacts the fish gut microbiome. This lack of insight is problematic given the frequency with which wild and managed fish are exposed to intestinal infections, as well as the potential for the gut microbiome to mediate these infections or their health impacts. In this study we report that infection with the nematode *P. tomentosa* results in rapid restructuring of the zebrafish gut microbiome. In addition, we find that the most dramatic disruption of the gut microbiome corresponds to time points with the greatest inflammation, and epithelial hyperplasia in the gut based on our parallel histology experiment. Finally, we find relationships between specific stages of the parasite life cycle and microbial abundance.

Our study design allowed us to determine how the gut microbiome changes over the course of infection by an intestinal parasite. Specifically, we exposed fish to the parasite by simulating their natural route of infection and tracked infection status and the gut microbiome over time. Prior work has established that infection with helminths can disrupt the structure of the microbiome [14]. For example, helminth infection has been linked to increased microbiome diversity in humans [10]. Others have shown drops in bacterial diversity during helminth infection in mammals [11,39]. Specific alterations in microbial taxonomic abundance after experimental infection with *Trichuris suis* [40], *T. muris* [39], and *H. polygyrus bakeri* [41] have also been reported. In the present study we found no evidence of altered microbial alpha-diversity, however, we do find changes in beta-diversity, indicating that infection with *P. tomentosa* results in a restructuring of the gut microbiome. We also observed a decrease in intra-individual beta diversity in all infected time points, except the terminal time point. Furthermore, we find evidence that the structure of the microbiome co-varies with the life history stage of the parasite in the intestinal tract. For example, significant differentiation in the beta-diversity of the microbiomes associated with the abundance of eggs, larvae, and adults. These results could indicate that *P. tomentosa* infection alters the host environment in such a way as to select for a specific conformation of the microbiome, and that the selection is differentially dependent upon the life history stage of the host. Alternatively, the bacteria in these *P. tomentosa* infected communities may simply be better able to survive in the inflammatory environment created by the parasite, and that the inflammatory context of the gut changes over the course of infection. We also cannot rule out the possibility that the microbiome of the parasite itself changes over time and influenced the alterations in microbial community composition and diversity observed in these experiments. It is clear however that *P. tomentosa* infection results in the rapid restructuring of the zebrafish gut microbiome and that the parasite’s life stage associates with gut microbiome composition.

Host-associated microbes can influence the colonization efficiency and pathogenicity of helminth parasites. For example, *Drosophilia neotestacea* harbors a maternally transmitted bacterium that protects the fly from the helminth parasite *Howardula aoronymphium* [42]. Similarly, *Lactobacillus casei*, and *Bifdobacterium animalis* reduce *Trichuris spiralis* [43] and *Strongyloides venezuelensis* [44] burden in mice, respectively. The microbiome is also necessary for some helminth infections. For example, *T. muris* requires the intestinal microbiome to establish infections in mice[45]. Similarly, more adult worms were recovered from mice with conventional microbiomes than germ-free mice when infected with *H. polygyrus bakeri* [46]. In the present study, we observed associations between worm burden and microbial abundance. For example, the abundance of several OTUs associated with the classes Gammaproteobacteria and Betaproteobacteria were positively correlated with the number of adult worms in the intestines of fish. Conversely, another group of OTUs associated with the classes Betaproteobacteria and Fusobacteria were negatively associated with adult burden. There are several possible explanations for these observations: 1) specific taxa promote or disrupt parasite colonization, growth and development [43–45], 2) some taxa may be better adapted to the altered gut microenvironment during infection and concordantly increase in abundance, 3) the parasite either opens niche space for microbial taxa to differentially colonized, or destroys niche space of resident bacteria, or 4) the composition of the parasite’s own microbiome varies over the course of infection. Further investigation is needed to determine if these correlations reflect causal relationships between microbial abundance in the gut microbiome and *P. tomentosa* infectivity or fecundity.

*Pseudocapillaria tomentosa* has been associated with zebrafish colony mortality and may be involved in the development of gastrointestinal tumors [17]. In contrast to previous reports wherein a subset of infected fish exhibit clinical pathologies [15,17], all exposed fish in the current study exhibited sub-clinical phenotypes. Here, pathological changes were confined to the epithelium and lamina propria, whereas in clinical zebrafish the worms often extend deeper in the intestinal lining and cause prominent coelomitis [17,47]. Mortality of the 5D line fish used in this study was also lower than previously reported in studies using the AB zebrafish line (2% vs ~16%) [24], possibly due to genotypic variation [48]. Interestingly, differences in microbiome composition have also been observed across different zebrafish strains [28], though it is unclear if genetic, environmental, microbial, or other factors contributed to the low mortality rates observed here. Importantly the inflammation and hyperplasia observed in this experiment may be an important factor in shifting microbiome structure during *P. tomentosa* infection. It is important to point out that the histological data and microbiome data were obtained from two separate cohorts of fish, therefore it is possible that the histological differences observed in this cohort were not precisely reflective of those we were unable to quantify in the other. Indeed, it seemed that the cohort used for histology progressed slightly faster than the fish used for the microbiome experiments (i.e., larvae present at 8d p.e. in the fish used for histology compared to no larvae 6d p.e. in the microbiome fish). We have shown in other studies that histology is more sensitive than wet mounts for small metazoan parasites of fish [49,50], and perhaps the larval population was not sufficiently large to be detected in the wet mount preparations. It is also possible that the slight difference in timing of these experiments might have led to this discrepancy. Future time course experiments exactly coupling fecal sampling and histological investigations would help to clarify how this perturbed gut microenvironment may impact microbiome structure during infection.

Infectious disease presents a major danger for maintaining functional experimental animal colonies. Not only do they pose risks to the health of the animals, which can disrupt research activities, but they can introduce potential confounding experimental results, especially in the case where the infection is cryptic [15]. Therefore, it is important to consider how colony infections may impact host physiology and the interpretations of experimental results. Infection of zebrafish with *P. tomentosa* resulted in restructuring of the microbiome that persisted over the duration of infection. Previous research has linked disruption of the microbiome with altered host physiology [31,51–53], behavior [54], and may contribute to the development and severity of disease [55,56]. As a result, it is concerning that this common infectious agent in zebrafish research facilities results in a significant perturbation to the gut microbiome, as there exists the potential that many experimental endpoints may be skewed as a result of an infection. An additional concern is that infection may yield a long-lasting impact on the operation of the gut microbiome, such that previously infected fish may not be appropriate for experimentation. Unfortunately, we did not follow fish to the resolution of infection, with the exception of a single individual, so it is difficult to determine the microbiome’s resiliency to infection. Future studies designed to follow individual fish across the length of infection and beyond are needed to determine the resiliency of the gut microbiome to helminth infection. Additionally, studies that quantify the specific impact of helminth-infection induced changes to the microbiome are needed.

These experiments demonstrate that infection with *P. tomentosa* alters the microbiome of zebrafish in a life stage dependent manner. Although it is unlikely that any specific results (i.e., altered taxa) can be broadly generalized to other intestinal helminth infections, we may be able to ultimately use the zebrafish to answer fundamental questions about how parasites, hosts and microbiome interact. This includes quantifying the resilience of the gut microbiome parasitic infection and determination of how the gut microbiome influences parasitic infection. Given that intestinal parasites exhibit a selective force on natural populations of animals[6], answering these questions may ultimately clarify one of the mechanisms through which the gut microbiome influences animal evolution. These studies would not only advance our understandings of host-parasite-microbiome interactions, but will also add significantly to our theoretical understanding of microbiome ecology.

## Acknowledgements

This study was supported in part by NIH ORIP 5 R24 OD010998 to M.L. Kent and National Council for Scientific and Technological Development (CNPq 202030/2014-8) grant to M.L. Martins. The authors thank the member of the CGRB for their assistance with sequencing and maintenance of our computational infrastructure.

---

**Supplemental Figure 1. Parasite burden kinetics.** Boxplots of **A)** larvae, **B)** egg, and **C)** adult parasite abundance across time points examined.

**Supplemental Figure 2. Microbiomes are more homogenous in fish with parasite burden.** Box plots of within-group Bray-Curtis dissimilarity.

